# *In vivo* bioreactor using cellulose membrane benefit engineering cartilage by improving the chondrogenesis and modulating the immune response

**DOI:** 10.1101/742759

**Authors:** Xue Guang Li, In-Su Park, Byung Hyune Choi, Ung-Jin Kim, Byoung-Hyun Min

**Author notes:** Corresponding author: Byoung-Hyun Min, M.D., Ph.D. Department of Orthopaedic Surgery, Ajou University School of Medicine, Wonchon-dong, Youngtong-gu, Suwon, Gyeonggi 443-749, Korea. Xue Guang Li and In-Su Park contributed equally to this work.

## Abstract

To regenerate tissue engineered cartilage as a source for the restoration of cartilage defects, we used a human fetal cartilage progenitor cell (hFCPC) pellet for improve the chondrogenesis and modulation the immune response with a *In vivo* (IV) bioreactor system, that was buried subcutaneously in the host and then implanted into a cartilage defect. *In vivo* bioreactor (IVB) was composed of silicone tube and cellulose nanopore-size membrane. FCPC pellets were first cultured *in vitro* for 3 days, and then cultured *in vitro*, subcutaneous and IV bioreactor for 3 weeks. First evaluated the IV bioreactor fluid appearance, component and liquidity, and then evaluate chondrogenesis and immunogenicity of the pellets using gross observation, cell viability, histology, biochemical analysis, RT-PCR, and Western Blot, finally evaluates the cartilage repair and synovial inflammation using histology. The fluid color and transparency of IV bioreactor were similar to synovial fluid (SF) and the component was also close to SF compared to the serum. IV bioreactor system not only promotes the synthesis of cartilage matrix and maintains cartilage phenotype, but also delays the occurrence of calcification compared with subcutaneous. A IV bioreactor, which has been predominantly adopted to study cell differentiation, was effective in preventing host immune rejection.

## 1. Introduction

Articular cartilage has a limited ability to heal after being damaged. As it has very little blood supply, cartilage has difficulty self-repairing after damage [1, 2]. Cell-based autologous or allogeneic transplantation has been employed to treat cartilage defects [3, 4]. The method of using autologous cells requires an invasive procedure for harvesting and has a limited amount of harvesting. Moreover, use of allogeneic MSCs has some limitations, an immune response may be elicited, leading to the failure of the cell therapy [5]. Furthermore, Chondrocytes dedifferentiate when expanding *in vitro*, resulting in loss of phenotypic and extracellular matrix (ECM) components [6].

Three factors play an important role in the engineering of cartilage tissue. Cells are somatic cells or stem cells [7]. Biomaterials are synthetic or natural materials. These elements are combined optimally and cultured in a bioreactor to produce cartilage similar to natural cartilage [8]. Tissue-engineered cartilage may be an applicable resource for healing cartilage defects, which would be more promising if the cartilage could be formed in an immunocompetent host [8]. However, inflammatory reactions, which might destroy chondrocytes and subsequently deform the tissue, obstructed reconstruction of tissue in immunocompetent animals. Moreover, host cell ingrowth would impair the quality of reconstructed tissue. Therefore, a new trend to avoid the above problems is *in vivo* bioreactor (IV bioreactor), which can effectively prevent vascular invasion, host cell invasion and immune rejection [7]. This using the body as a bioreactor, the traditional three elements (scaffold, cell and growth factor) were cultured, and the regeneration environment of the body provided help for its growth. This would be applicable for allografts because the immune system cannot recognize allografts once host cell invasion is inhibited. Cellulose is a kind of bio-inert material with good biocompatibility and low immune response, which is often used in medical implants [9]. The cellulose membrane of IV bioreactor used in this study is composed of fairly long cellulose fibrils connected with each other and has a highly porous three-dimensional network [10]. On the other hand, the selection of favorable stem cells in cartilage tissue engineering is of great importance. In previous stem cell treatments, stem cells not only have the ability for regeneration in chronic tissue damage but also have a regulatory effect on the immune environment. Implanted stem cells can regulate the immune environment during tissue repair for tissue regeneration [11]. Previously, we have reported that human fetal cartilage progenitor cells (hFCPCs) having high yield, proliferation, multipotent differentiation and maintains the chondrogenic phenotype abilities, in cartilage tissue formation [11]. Pellet culture was widely used in chondrocytes culture [12]. However, the chondrocyte pellet culture method can easily cause cartilage mineralization *in vitro* [13]. The subcutaneous environment was often used to the animal model of regenerate ectopic tissue-engineered cartilage, but due to the cell ingrowth and vascular invasion of the host, lead to a matrix is destroyed and calcified [8].

In this study, component and liquidity were analyzed of the cellulose membrane IV bioreactor fluid and evaluated the chondrogenesis, immunogenicity and the ability to healing cartilage defects of FCPCs pellets cultured in the bioreactor and compared with *in vitro* and subcutaneous.

## 2. Materials and Methods

### 2.1. Cell isolation and culture

Human fetal cartilage progenitor cell (hFCPC) were isolated from the human fetal cartilage tissue, as previously described [11]. The study was approved by the institutional review board (IRB) of the Ajou University Medical Center (AJIRB-CRO-07-139) and was carried out with the informed consent of all donors. All experiments were performed in accordance with relevant guidelines and regulations. Human fetal cartilage tissues (*n* = 2, F12w-c, M11w) were obtained from patients following elective termination at 12 weeks after gestation, and cells were isolated from the femoral head of the cartilage tissue. Cartilage pieces were digested in 0.1% collagenase type II (Worthington Biochemical Corp, Freehold, NJ, USA) in high-glucose Dulbecco’s modified Eagle Medium (DMEM; HyClone, Logan, UT, USA) containing 1% fetal bovine serum (FBS; HyClone) at 37 °C under 5% CO2. After 16 h, isolated cells were cultured in DMEM supplemented with 1% antibiotic-antimycotic and 10% FBS at a density of 8 × 10^3^ cells/cm^2^. After 3 days, non-adherent cells were removed and the medium was changed. Cells were passaged at 80% confluence by 0.05% Trypsin–EDTA (Gibco, Gaithersburg, MD, USA) treatment.

### 2.2. Pellet culture

Aliquots of 3 × 10^5^ cells/0.5 ml were centrifuged at 500 g for 10 min in 15 ml polypropylene tubes. After 1 day, cell pellets were cultured in chondrogenic defined medium (DMEM supplemented with ITS mixture, 50 μg/ml ascorbate-2 phosphate, 100 nM dexamethasone, 40 μg/ml proline, and 1.25 mg/ml BSA) without TGF-β, a typical chondrogenic inducer. Pellets were cultured for 3 days.

### 2.3. Preparation of cellulose membrane chamber

For this study, our own cellulose membrane chamber was manufactured. Cellulose membrane chamber was composed of a silicone tube (21 mm × 15 mm × 10 mm, outer diameter × inner diameter × height) (Korea Ace Scientific, Seoul, Korea) and cellulose membrane. The pore size of the cellulose membrane is about 50-400 nm (supporting information Figure 1), provided by the bioenergy research center, college of life sciences, Kyung Hee University [10]. Both ends of the silicone tube were sealed with a cellulose membrane and fixed using 7-0 black silk. Followed by sterilization in an autoclave at 121 °C for around 20 minutes.

### 2.4. Ectopic chondrogenesis in the subcutaneous and IV bioreactor environment

This study was approved by the ethics committee for animal research of the Laboratory Animal Research Center of Ajou University Medical Center (approval No. 2014-0068). Authors complied with institutional ethical use protocols (NIH Guide for Care and Use of Laboratory Animals). 3.5 kg female adult New Zealand white rabbits (*n* = 24 for subcutaneous and IV bioreactor groups; OrientBio, Seongnam, Korea) were anesthetized with a mixture of Zoletil and Rumpun. manufactured cellulose membrane chambers (two chambers/rabbit) were implanted under back skins of rabbits. In this study total of 360 pellets (pellet size 1.14 ± 0.04 mm) were divided into three environments: *in vitro*, rabbit subcutaneous and IV bioreactor. After 3 days, a total of 240 pellets were put into the subcutaneous and cellulose membrane chamber (5 pellets/chambers). Rabbits were sacrificed at each time point of 1 day, 1, and 3 weeks post-implantation. Retrieved pellets were used for analyses, gross observation, pellets size measurement, cell viability assay, histological analysis, immunohistochemical analysis, biochemical, reverse transcription polymerase chain reaction (RT-PCR) analysis and western blot analysis, respectively.

### 2.5. Cartilage defect repair

The same 45 adult New Zealand White rabbits were used for cartilage defect repair (OrientBio). Under general anesthetic, the knee was exposed after lateral skin incision and muscle detachment. A cylindrical cartilage defect (diameter: 2.0 mm, depth: 0.5 mm) was created in the trochlear groove of the femur. Then, 1 week cultured engineered cartilage tissues from *in vitro*, autologous IV bioreactor and subcutaneous, were implanted into the defects and the wound was stitched layer to layer. The control group was implanted with nothing at the defect. After surgery, all of the rabbits were kept in individual cages at constant temperature and humidity, with unrestricted access to a standard diet and water. Rabbits could walk freely with full weight bearing. Both analgesics and antibiotics were administered for 3 days after surgery. The animals were sacrificed using an intravenous injection of a euthanasia solution at 4, 8 and 12 weeks after surgery, then the repaired cartilage and synovial membrane was harvested for gross observation, H&E staining, Safranin-O staining and Immunohistochemistry (IHC) staining of collagen type II (COL II), collagen type I (COL I) and collagen type Ⅹ (COL Ⅹ).

### 2.6. Measurement of transmittance

Initially, the transparency of each fluid was visually estimated. Subsequently, the transmission spectra of each fluid were observed using a UV-Vis spectrophotometer (Jasco V-650, Japan). The spectral distribution was measured in the visible wavelength range (400-800 nm). The results were normalized with water.

### 2.7. Permeability assay

In order to test diffusion across the cellulose membrane, we confirmed the diffusion patterns of glucose through the cellulose membrane using a two-chamber system developed previously in our laboratory [14]. Briefly, the cellulose membrane was fixed between two specific chambers. We loaded 4500 mg/L glucose DMEM on one side of the chamber and loaded distilled water (DW) on the other side of the chamber. After 2, 4, 8, 12, and 24 hours measured the glucose concentration in the DW in the Ajou University Hospital.

### 2.8. In vivo bioreactor fluid component analysis

Total protein and glucose in IV BIOREATOR fluid were measured at the Ajou University Hospital. Hyaluronan (HA) content in IV BIOREATOR fluid was measured using ELISA kit according to the manufacture’s instructions (R&D Systems, Minneapolis, USA). Lactate content in IV Bioreactor fluids was measured using EnzyChromTM L-Lactate Assay Kit (ECLE-100) according to the manufacturer’s instructions (BioAssay Systems, Hayward, CA, USA). Experiments were conducted in triplicate, and optical densities were used to normalize the lactate production results.

### 2.9. Gross observation and size measurement of the pellets

Retrieved pellets (*n* = 5/time point/group) were observed in terms of their shape, color, and size. The size of the pellets was determined using the Image J program.

### 2.10. Cell viability assay

The pellets (*n* = 5/time point/group) were incubation at 48-well plate in serum-free DMEM medium 200 μl alamarBlue (Invitrogen, California, USA) 20 μl, 37 °C in a 5% CO_2_ incubator. After 5h of incubation, the absorbance of duplicate samples (100μl) of each well was measured in a 96well plate, at wavelengths of 570 and 600 nm, using a microplate reader (Infinite M200, Tecan, CH).

### 2.11. Histology and Immunohistochemistry

Samples (*n* = 5/time point/group) were fixed with 4% formaldehyde for 24 h, dehydrated, and then embedded in paraffin wax. Sections, each 4 μm in thickness, were prepared and stained with hematoxylin/eosin for visualizing cell distribution and morphology (for assessment of synovitis using Krenn’s synovitis score system [15], supporting information Table 1), Safranin-O/fast green for accumulation of sulfated proteoglycans (for the evaluation of Safranin O-Fast green stained cartilaginous pellet using the Bern score [16], supporting information Table 2 and cartilage repair score using the O’Driscoll score system [17], supporting information Table 3), and Alizarin Red for calcified pellets (for quantitative analysis of the calcification area using the Image J program). For immunohistochemical analyses, sections of samples prepared as described above were treated with 3% Hydrogen peroxide (H_2_O_2_) for 10 min and reacted with a pepsin solution (Golden Bridge International, Inc, Mukilteo, WA, USA) for 10 min. After blocking the sections with 1% BSA in phosphate-buffered saline (PBS) for 1 h, sections were incubated with primary antibodies (1:200) for 1.5 h at room temperature. The primary antibodies used were type II, type I and type X collagen (all from Abcam, Cambridge, UK; supporting information Table 4). With 2 times washing in PBS, sections were incubated with a biotinylated secondary antibody (1:200) for 30 min and peroxidase-conjugated streptavidin solution for 30 min at room temperature (both from Golden Bridge International). Finally, sections were reacted with a 3,3’ -diaminobenzidine (DAB) solution (Golden Bridge International) and counterstained with Mayer’s hematoxylin (Sigma, St Louis, MO) and then mounted for microscopic observation (Nikon E600, Japan).

### 2.12. Biochemical assay

For DNA and sulfated glycosaminoglycan (sGAG) contents assays, pellets (*n* = 5/time point/group) were fully digested for 24 h at 60 °C in papain digestion solution containing 125 μg/ml of papain with 5 mM L-cysteine-HCl and 5 mM EDTA in 100 mM Na_2_HPO_4_ (all from Sigma, USA). The DNA content was measured using the Picogreen assay. Total sGAG content was measured using the Blyscan GAG assay kit (Biocolor, Carrickfergus, UK). The papain-digested pellets were reacted with Blyscan dye reagent for 30 min and centrifuged for 10 min (12,000 rpm). The deposits were dissolved with dissociation reagent and absorbance was read at 656 nm. For collagen content assay, pellets (*n* = 5/time point/group) were fully digested for 2 days at 4 °C in 0.1M of HCl digestion solution containing 1 mg/ml pepsin (Sigma). Collagen content was measured using the Sircol collagen assay kit (Biocolor). The HCL-digested pellets were reacted with Sircol dye reagent for 30 min and centrifuged for 10 min (12,000 rpm). The deposits were washed with ice-cold acid-salt wash reagent and centrifuged for 10 min (12,000 rpm). The deposits were dissolved with Alkali reagent and absorbance was read at 555 nm.

### 2.13. Reverse Transcription Polymerase Chain Reaction (RT-PCR)

The pellets (*n* = 5/time point/group) were rinsed in PBS and lysed in 500 μl TRIzol (Invitrogen, CA, USA) for 10 min. Total RNA was extracted according to the manufacturer’s instructions. Then, cDNA was synthesized using total RNA and iScript cDNA synthesis kit (Bio-Rad, Hercules, CA, USA). Reverse transcriptase-polymerase chain reaction (RT-PCR) was performed for human leukocyte antigens-ABC (HLA-ABC), CD80, CD86 and glyceraldehyde-6-phosphate dehydrogenase (GAPDH) using AccuPower PCR PreMix (Bioneer, Korea) according to the manufacturer’s instructions. The sequences of the primers and the reaction conditions are described with detailed information in Table 1.

**Table 1.**
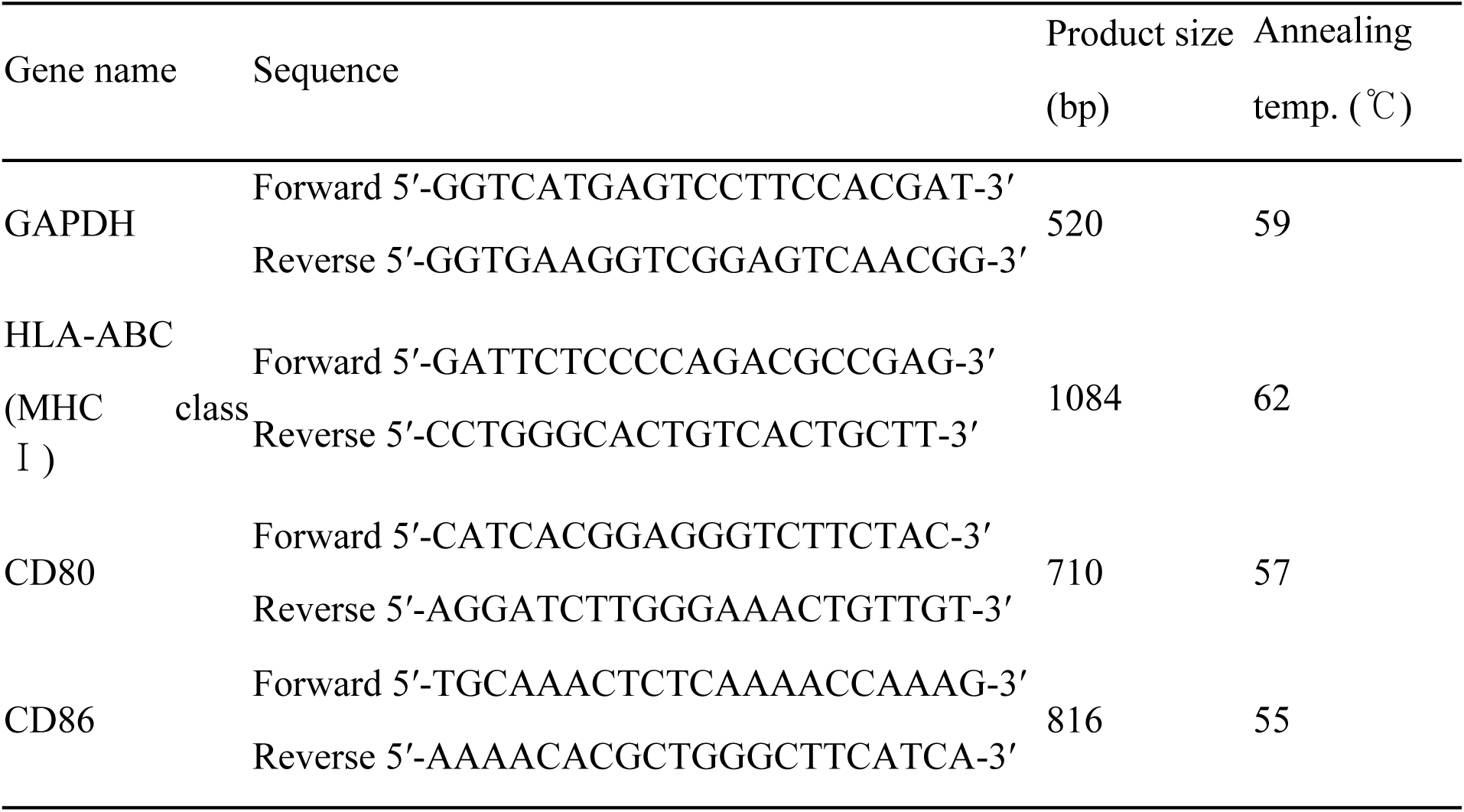
Sequences of primers used for RT-PCR analysis. The expected size of products and annealing temperatures of the PCR reactions are also presented.

### 2.14. Western Blot analysis

Western blotting for HLA-ABC, CD80, CD86, and β-Actin. Total proteins were extracted from pellets and using Radioimmunoprecipitation assay (RIPA) lysis buffer (Rockland, Gilbertsville, PA, USA) and the protein concentrations were quantified using the Bradford assay (Bio-Rad). The protein extracts were subjected to electrophoresis through a 4–20% precast polyacrylamide gel (Bio-Rad). Subsequently, the proteins were transferred onto polyvinylidene fluoride (PVDF) membranes (Bio-Rad) and the membranes were blocked in 5% nonfat dry milk in Tris-buffered saline (TBS) containing 0.1% Tween-20 (TBST). Membranes were incubated for 2 h at 37 °C with primary antibodies against β-Actin (GeneTex, Irvine, CA, USA; dilution of 1:2000), HLA-ABC, CD80, and CD86 (all from Abcam; dilution of 1:2000; supporting information Table 4). The membranes were then incubated with horseradish peroxidase (HRP)-conjugated anti-rabbit or anti-mouse secondary antibodies (both from GeneTex; dilution of 1:2000) at 37 °C for 1 h.

### 2.15. Statistical analysis

Data were analyzed for statistical significance by one-way and two-way analysis of variance (ANOVA) Tukey’s multiple comparisons test using GraphPad Prism 7.00 software. The experiments were repeated at least three times (*n* = 5). Data are expressed as mean ± standard deviation (SD). Statistical significance was assigned as \**p* < 0.05.

## 3. Results

### 3.1. Set up the in vivo bioreactor

In order to confirm own self-regenerative capacity to regenerate new cartilage, we created the *in vivo* bioreactor using cellulose membrane and silicon tube in rabbit subcutaneous (Fig. 1A-D). The FCPC pellet culture within the chamber at 3 days after implantation, a thin fibrous tissue encapsulated with a vascular network surrounded all chambers, and the only minimal fibrotic reaction was seen in the subcutaneous pockets surgically created. The body fluid was full inside the IV Bioreactor chamber. (Fig. 1E). The pellet culture 1 week after, it was transplanted autologous cartilage defect (Fig. 1F). Pellet was opaque and whitish, like native cartilage, and no vascular invasion was observed. The shape of the newly formed tissue changed little compared with that before implantation.

**Figure 1.**
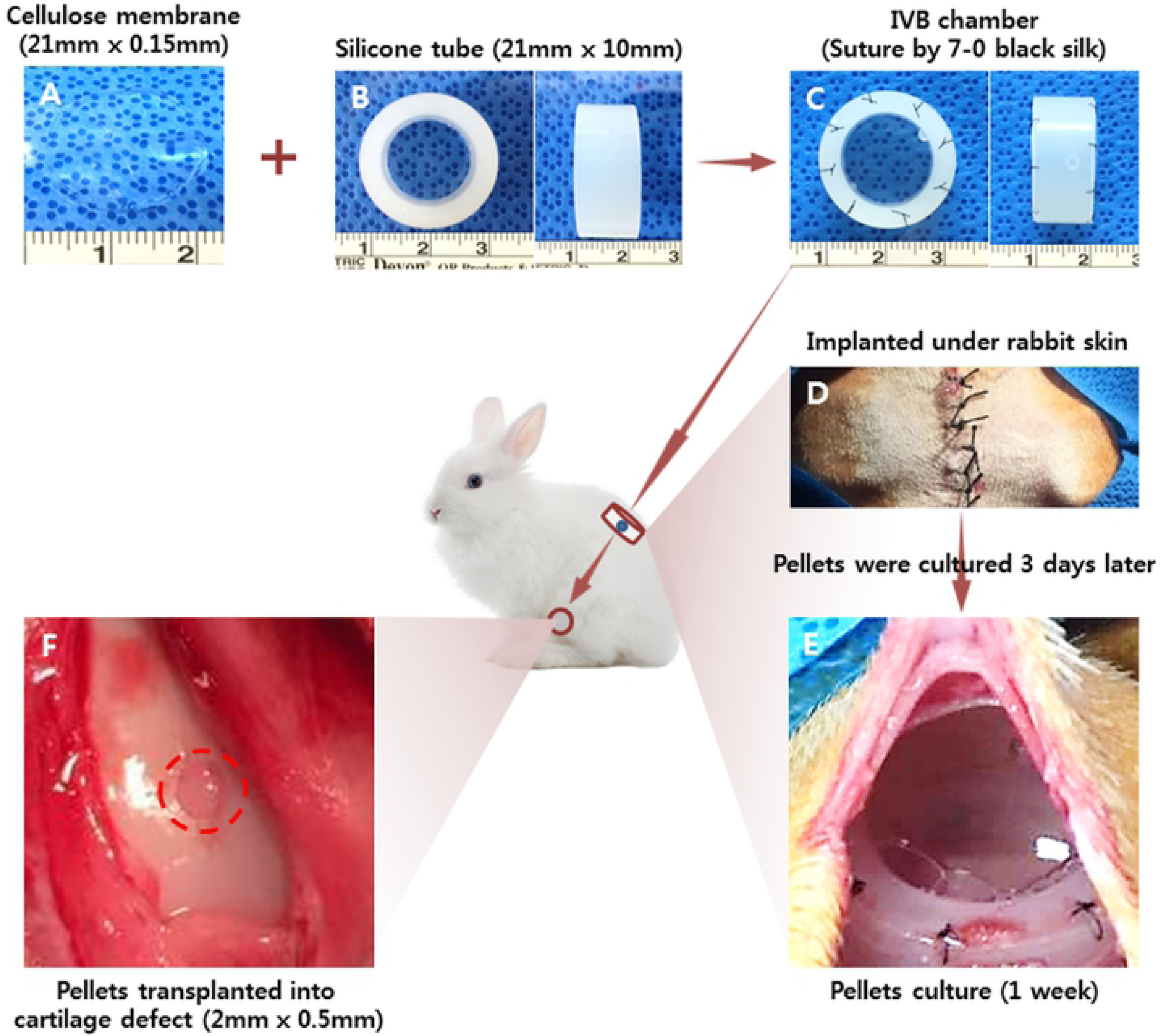
Schematic illustration of the overall design of this study. FCPC pellets were used to construct ectopic cartilage in the IV BIOREATOR system for cartilage repair. (A) A gross image of the cellulose membrane (size 21 mm × 0.15 mm). (B) A gross image of the silicone tube (size 21 mm × 10 mm). (C) A gross image of the IV bioreactor chamber suture by 7-0 black silk. (D) A gross image of the two IV bioreactor chamber implanted under the skin of a rabbit. (E) 3 days after IV bioreactor chamber implantation, pellets were placed into the IV bioreactor for culture, and the gross image of IV bioreactor was obtained after 1 week of pellets culture. (F) A gross image of the 1 week culture of pellets transplanted into the cartilage defect (defect size 2 mm × 0.5 mm).

### 3.2. IV BIOREATOR fluid characteristics and Confirm the fluid liquidity

The appearance (Fig. 2A) and transparency (Fig. 2B) of the IV bioreactor fluid and native synovial fluid (SF) are shown. IV Bioreactor fluid recovered at after pellets culture 1day, 1 and 3 weeks. The control was a failed cellulose membrane chamber that was yellow-brown, turbid and contained small particles, but the IV Bioreactor fluid is clear, slightly yellow and free of particulate, at all time points, similar to SF. The light transmittance results showed that the synovial fluid was the highest, followed by the IV Bioreactor fluid at 1 day, 1 and 3 weeks, and the lowest was the control. The transmittance was 97%, 96%, 94%, 89% and 63% respectively at 800 nm. The contents of total protein, glucose, and hyaluronic acid in IV Bioreactor fluid before pellets culture were measured and compared with those in rabbit SF and serum, and the contents of human SF and serum were referenced (Table 2). The results showed that these components in the IV BIOREATOR group were closer to synovial fluid than serum. Before and implantation 3 weeks after the permeability of the cellulose membrane used in the IV BIOREATOR was compared for 24 hours (Fig. 2C). The color thickens with the time of the right chamber (DW) and the color of both chambers has changed similarly at 24 hours in both groups. The concentration of the glucose in the right chamber was similarly increased with time at both groups (Fig. 2D). Increased quickly from the first 2 hours and relatively slowly since then. Total protein, glucose and lactic acid contents in IV BIOREATOR fluids with and without pellets were compared at 3, 6, 9 and 12 days (Table 3). The contents of these components were similar between both groups at the same time point, and each group maintained similar levels at all time points. These results indicated that the IV BIOREATOR fluid content was closer to SF than serum. And after 3 weeks of implantation, the cellulose membrane can be seen that the permeability is good as before. And the IV BIOREATOR system maintained homeostasis.

**Figure 2.**
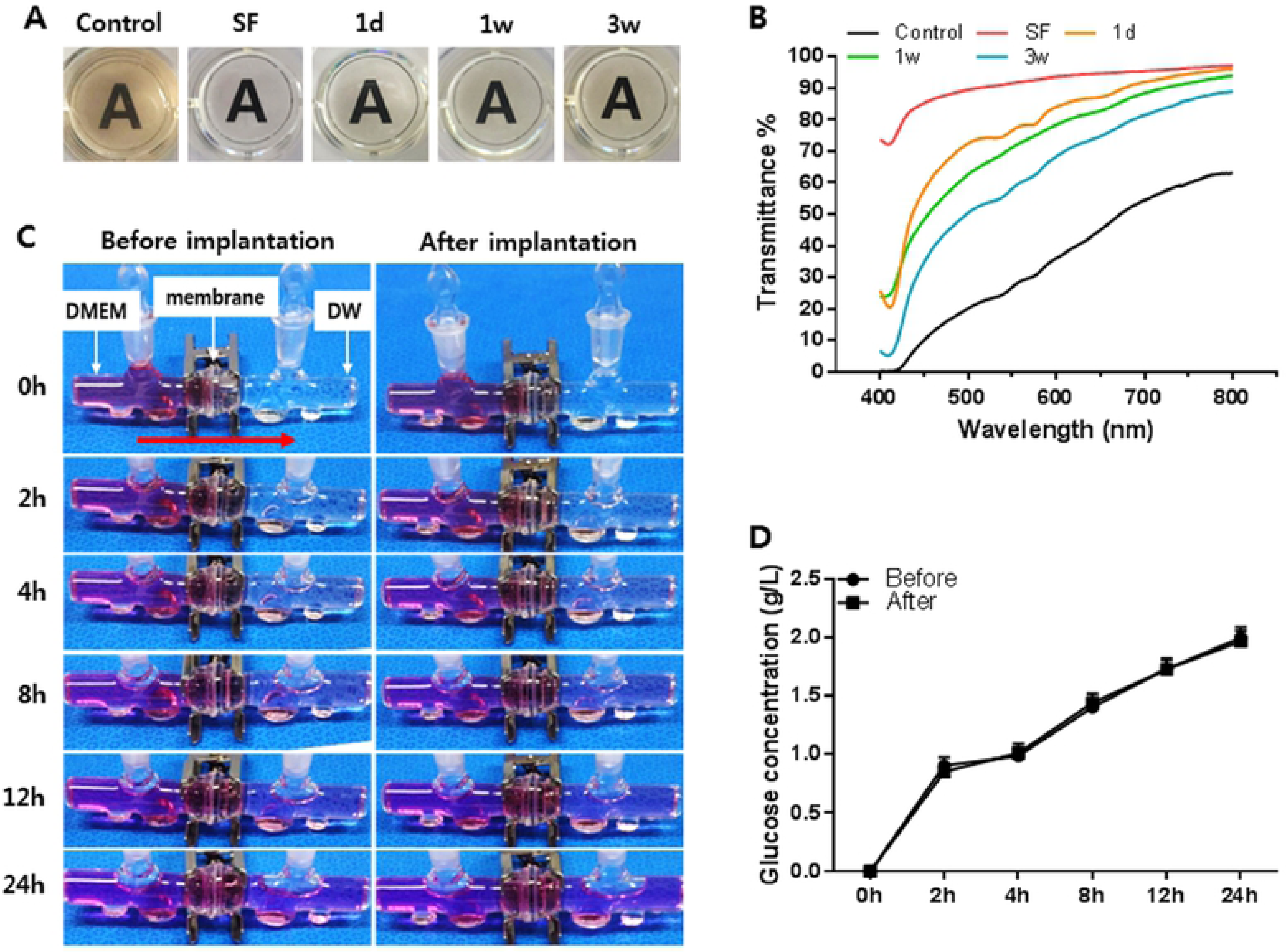
IV bioreactor fluid characteristics and Confirm the fluid liquidity. The IV bioreactor fluid appearance (A) and transparency (B). (C) The permeability of the cellulose membrane. (D) The concentration of glucose that was transferred from the left chamber to the right through the cellulose membrane.

**Table 2.** IVB fluid component analysis

|  | Rabbit |  |  | Human * |  |
| --- | --- | --- | --- | --- | --- |
|  | Serum | SF | IVB fluid | Serum | SF |
| Total protein (mg/ml) | 63 ± 3.9 | 25.5 ± 3.2 | 35.2 ± 3.1 | 67.2 ± 3.1 | 19.9 ± 5 |
| Glucose (mg/dl) | 114 ± 16 | 97 ± 9 | 106 ± 12 | 108 ± 18 | 93 ± 17 |
| Hyaluronan (ug/ml) | 0.2 ± | 30 ± 2.9 | 5.1 ± 0.3 | 0.3 ± | 2500 ± 790 |

\* Source: [25] Balazs EA, et al. Arthritis Rheum. 1967;10(4):357–76.

**Table 3.**
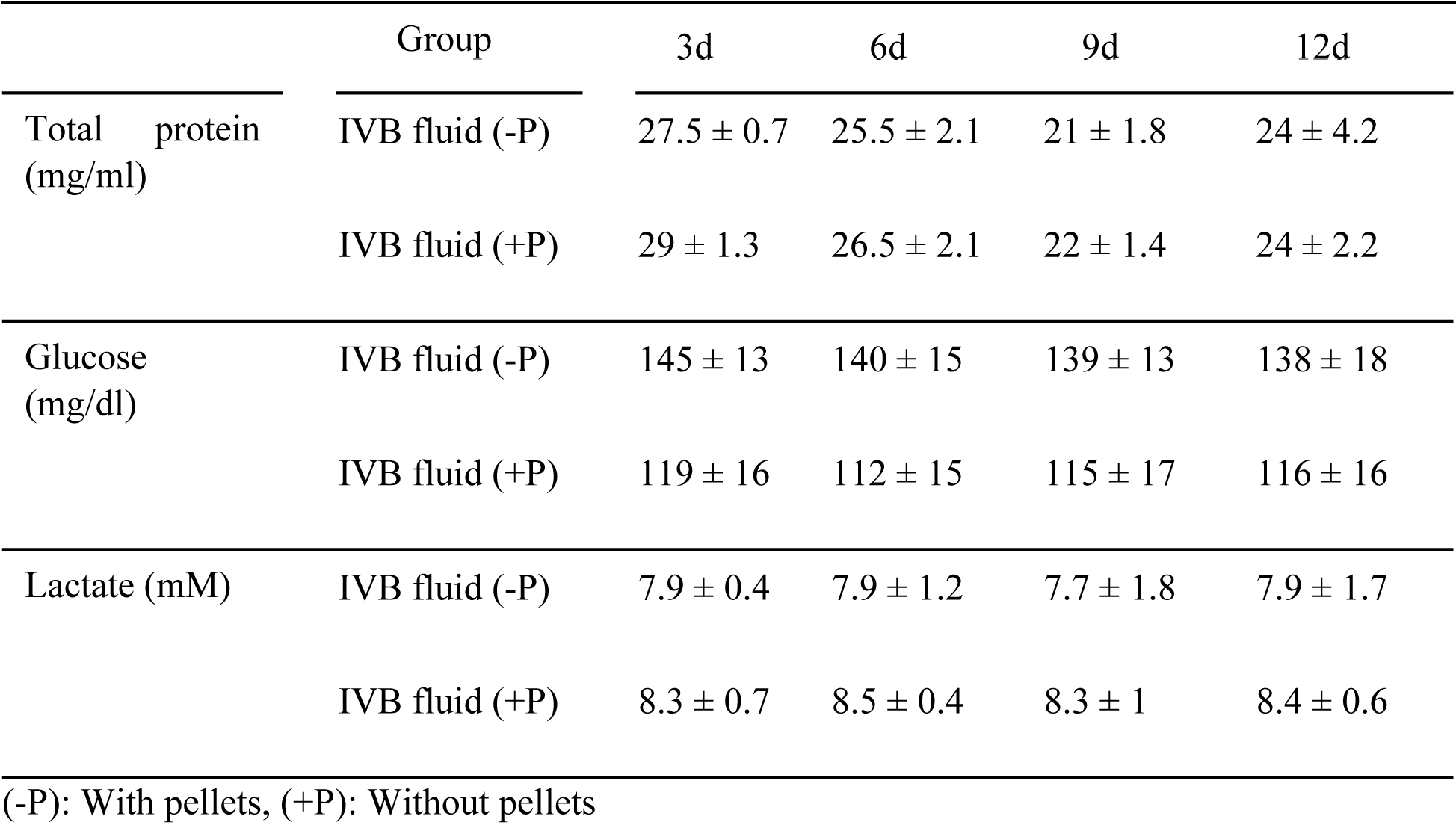
Confirm the IVB fluid liquidity

### 3.3. Gross observation, size measurement and cell viability of the pellets

The effect of the IV bioreactor was compared with *in vitro* and subcutaneous on the growth of FCPC pellets. The pellets were first cultured *in vitro* for 3 days, then cultured *in vitro*, subcutaneous and IV Bioreactor for 1day, 1 and 3weeks. All groups pellets showed whitish, hyaline cartilage-like circular morphology and subcutaneous tissue adhesion or turned a blood stained red color did not appear at any time point (some subcutaneous groups showed adhesions, data not shown) (Fig. 3A). In the gross images and size measurement, the size of the *in vitro* and IV bioreactor groups significantly increased with time, but the subcutaneous group was decreased at 3 weeks. The estimated size of the IV Bioreactor group (1.762 ± 0.132 mm^2^) was larger than that of the subcutaneous group (1.244 ± 0.115 mm^2^) (*p* < 0.05), and similar to *in vitro* group (1.969 ± 0.135 mm^2^) at 3 weeks (Fig. 3B). The pellet viability was measurement (Fig. 3C). The reagent changed from purple to pink of all groups at all time points and measured the absorbance (Fig. 3D). The absorbance increased with time at *in vitro* and IV Bioreactor groups, but the subcutaneous group was decreased at 3 weeks. These results indicated the ability of the IV bioreactor to maintain the growth of FCPC pellets observed above by gross and viability analysis.

**Figure 3.**
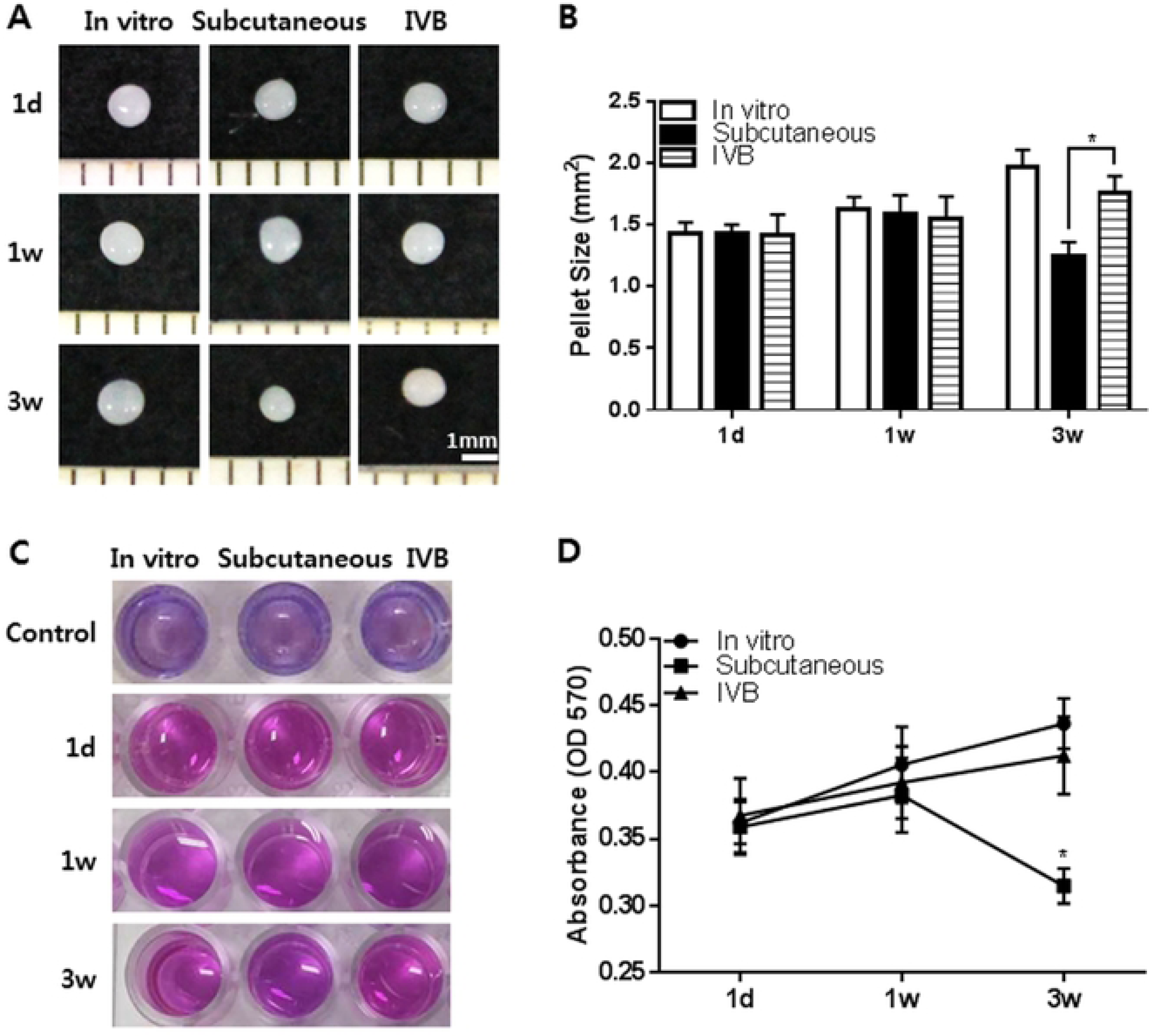
Gross observation, size measurement and cell viability of pellets retrieved from the *in vitro*, subcutaneous and IV bioreactor experiment. The pellets were cultured for 3 days *in vitro* before implantation in the subcutaneous and IV bioreactor of rabbit. The implanted pellets (*n* = 5/time point/group) were retrieved at 1day, 1 and 3 weeks after implantation. (A) The gross images of pellets are presented. (B) The size of each pellet was measured in the *in vitro* (open bar), subcutaneous (solid bar) and IV bioreactor (horizontal striped bar) groups using Image J program. (C) The cell viability was examined using Alamar blue assay. (D) Absorbance of Alamar blue at 570 nm. The data are presented as a mean ± standard deviation (SD) from 5 independent experiments (*n* = 5). \**p* < 0.05.

### 3.4. Histological observation of the pellets

Chondrogenesis of the pellets cultured *in vitro*, subcutaneous and IV bioreactor was further examined by histological analysis (Fig. 4A-B). Safranin-O/fast green staining showed that the accumulation of sulfated GAGs was significantly increased with time and no significant difference was noted in the intensity and area of Safranin-O staining between the *in vitro* and IV bioreactor groups until 3 weeks, but the subcutaneous group was decreased at 3weeks. The Bern score of the IV bioreactor group (5.8 ± 0.9) was larger than that of the subcutaneous group (4.0 ± 0.8) (*p* < 0.05), and similar to *in vitro* group (6.5 ± 1.2) at 3 weeks (Fig. 4B). In the high magnification images (x 400), the differentiated cells in the metachromatically stained area of the *in vitro* and IV bioreactor groups were round or long fusiform during the entire culture process, and lacunar structure characteristic of native cartilage is not seen (Fig. 4A-B). This morphology was observed for differentiated cells of the subcutaneous group in 1 week of culture, but the cells structures were severely degraded in the 3 weeks of culture.

**Figure 4.**
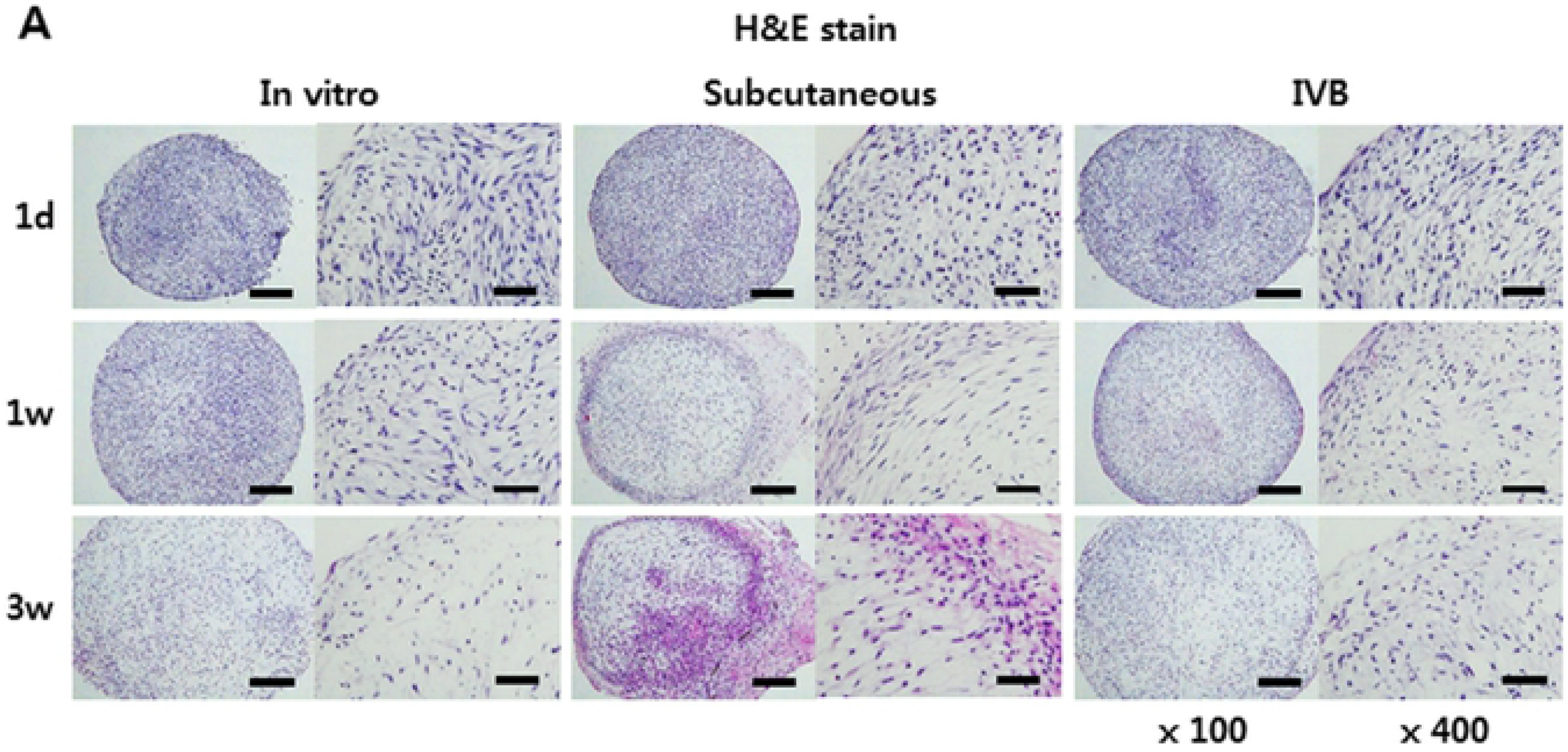
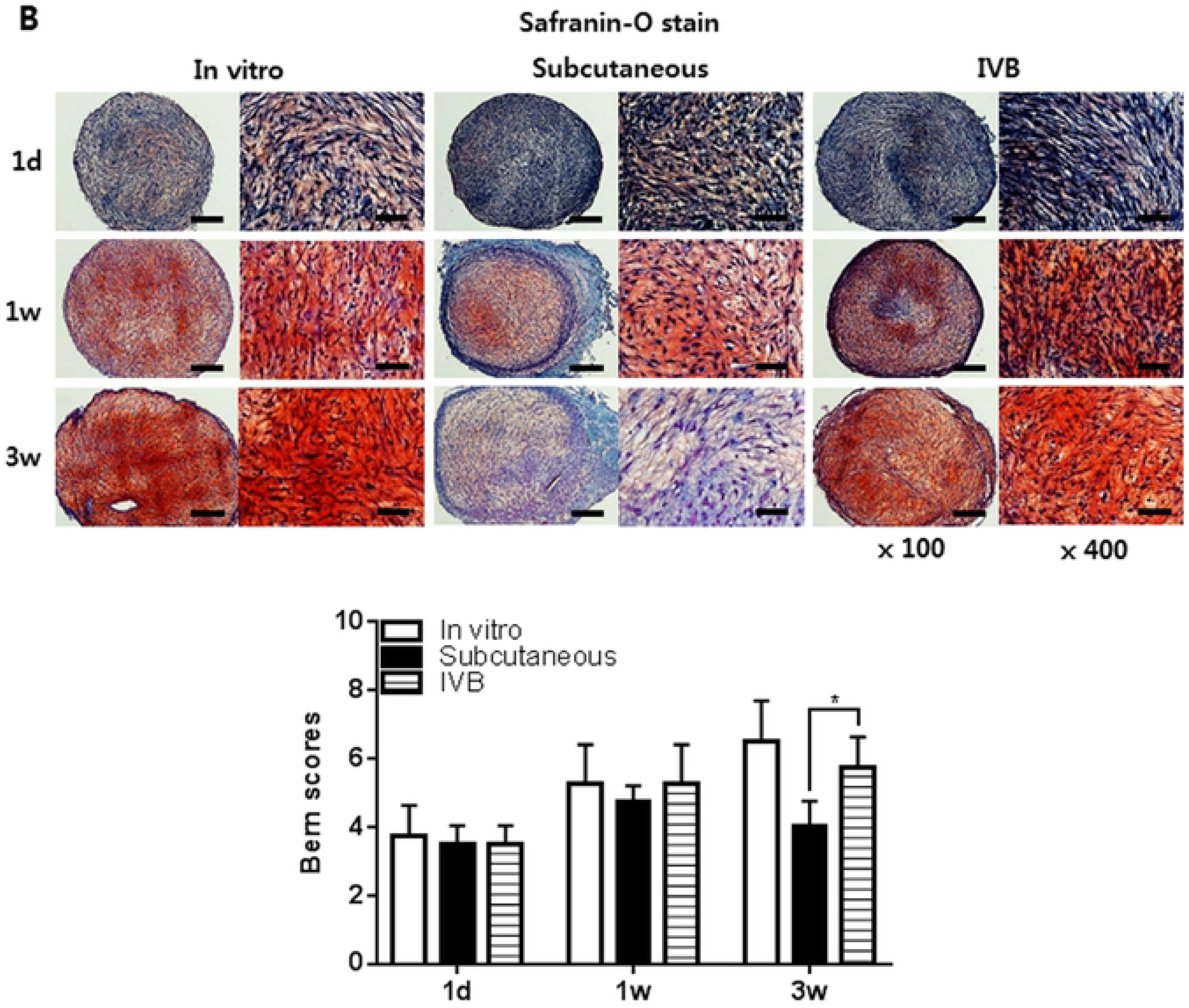
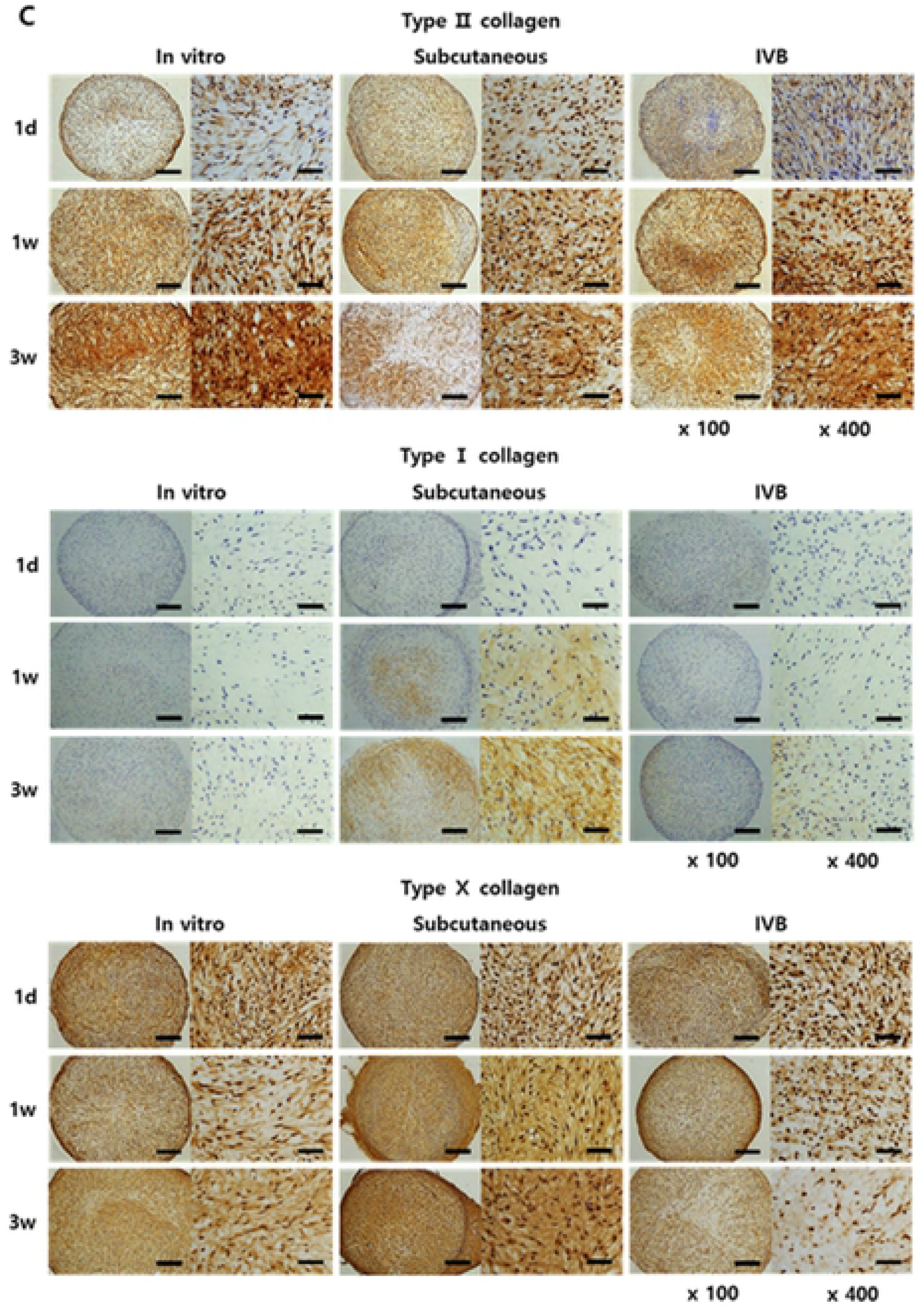
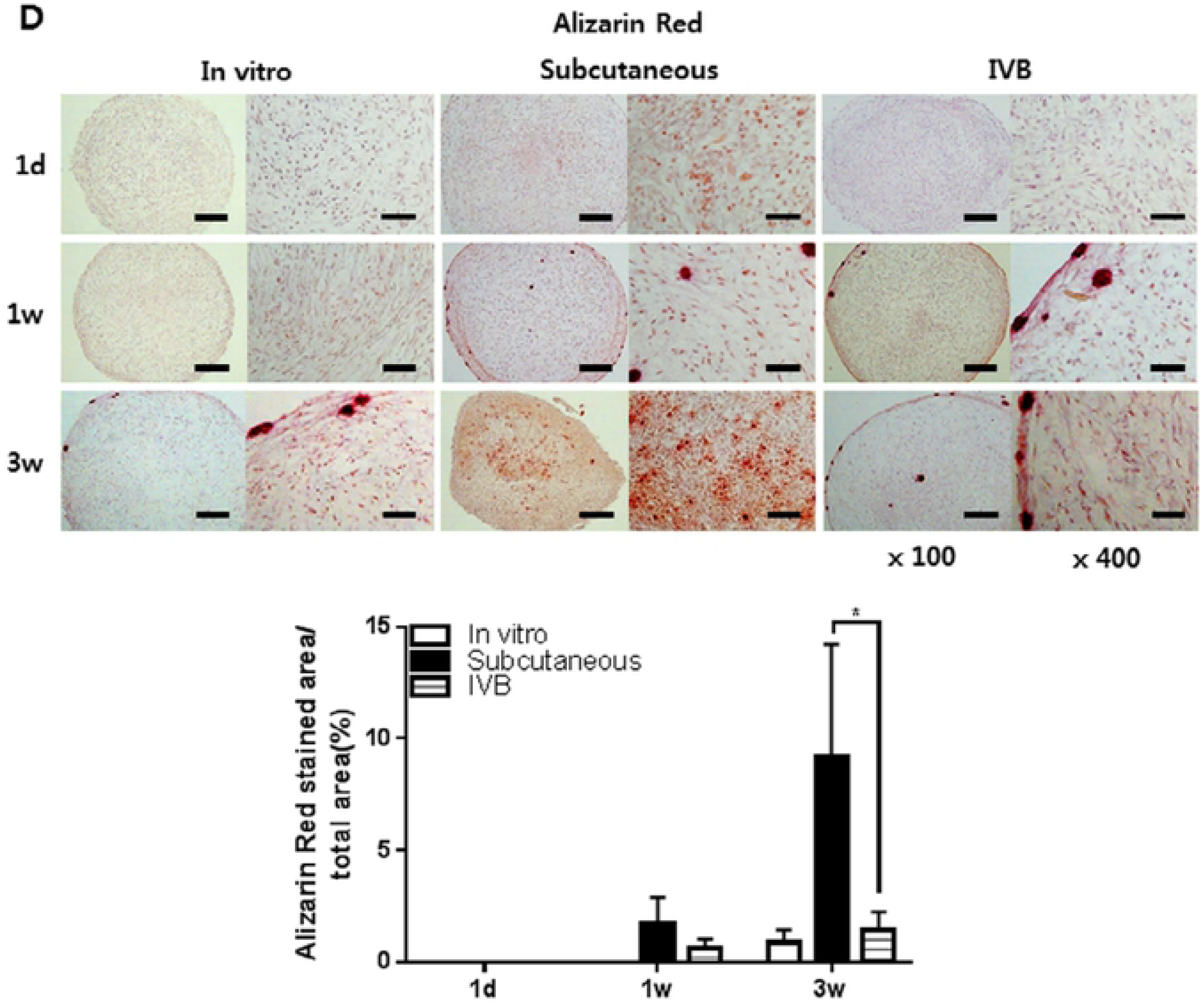
Histological and immunohistochemical observation of the pellets. The pellets were cultured for 3 days *in vitro* before implantation in the subcutaneous and IV BIOREATOR of rabbit. The implanted pellets (*n* = 5/time point/group) were retrieved at 1day, 1 and 3 weeks after implantation. The pellets were processed to prepare thin sections, each 4 μm in thickness. (A) Hematoxylin/eosin to observe the distribution of cells. (B) Safranin-O/fast green to observe accumulation of sulfated proteoglycans, and the Bern scores were evaluated. (C) Immunostained with an antibody against rabbit type Ⅱ, type Ⅰ and type Ⅹ collagen. (D) Alizarin Red stain to observe calcium deposits in the pellets. The stained images are presented as a whole pellet (left columns, × 100) and at high magnification (right columns, × 400). Scale bar = 200 μm for × 100 and 50 μm for × 400 images. Data are presented as a mean ± SD from 5 independent experiments in the histograms (*n* = 5). \**p* < 0.05.

### 3.5. Immunohistochemical observation of the pellets

The immunostaining for type Ⅱ, type Ⅰ, and type Ⅹ collagen further examined to confirm the chondrogenic phenotypes of the pellets (Fig. 4C). The type Ⅱ collagen increased with time *in vitro* and IV bioreactor groups, but in a subcutaneous group it decreased, reaching about 50% at 3 weeks in both its intensity and stained area than 1 week. The IV bioreactor group showed strong expression of type Ⅱ collagen in the whole pellets at 3 weeks and stronger than the subcutaneous group, but weaker than *in vitro* group. The type Ⅰ and type Ⅹ collagen showed weak expression at all time points *in vitro* and IV bioreactor groups, but in a subcutaneous group, it increased with time.

### 3.6. Calcification of the pellets

To examine if the loss of chondrogenic phenotypes was correlated with calcification of the matrix, Alizarin Red staining was performed, and using the Image J program confirmed that the percentages of calcification area (Fig. 4D). In both the subcutaneous and IV bioreactor groups, red stains, indicative of calcified mineral deposits, was first observed in the peripheral region at 1 week and spread into the central region at 3 weeks. The *in vitro* group it showed weak stain in the peripheral region at 3 weeks. The stained area was more intense and broader in the subcutaneous group (9.17 ± 5.04%) than in the IV bioreactor (1.43 ± 0.83%) (*p* < 0.05) and *in vitro* groups (0.88 ± 0.57%) (*p* < 0.05) at 3 weeks. These results indicated that the IV BIOREATOR not only better supports the chondrogenesis but also maintain the chondrogenic phenotype of FCPC pellets.

### 3.7. Biochemical analysis for the content of DNA, GAGs and collagen

The content of DNA, GAGs, and collagen was measured quantitatively by chemical assays of the retrieved pellets (Fig. 5). The DNA content increased with time after culture *in vitro* and IV bioreactor groups, but subcutaneous group was decreased at 3 weeks (Fig. 5A). The DNA content of the IV bioreactor group (1.40 ± 0.11 μg/mg) was higher than of the subcutaneous group (1.08 ± 0.08 μg/mg) (*p* < 0.05), but lower than that of the *in vitro* group (1.56 ± 0.07 μg/mg) (*p* < 0.05) at 3 weeks. This result suggests that the number of cells increased with time *in vitro* and IV Bioreactor groups, but the subcutaneous group was decreased at 3 weeks. The content of GAGs and collagen showed a similar pattern overall to that of DNA content. The GAGs content increased with time after culture *in vitro* and IV bioreactor groups, but the subcutaneous group was decreased at 3 weeks (Fig. 5B). The IV bioreactor group (6.15 ± 0.27 μg/mg) was higher than of the subcutaneous group (2.79 ± 0.35 μg/mg) (*p* < 0.05), and similar to *in vitro* group (6.98 ± 0.26 μg/mg) (*p* < 0.05) at 3 weeks. The collagen content increased with time after culture *in vitro* and IV bioreactor groups, but the subcutaneous group was decreased at 3 weeks (Fig. 5C). The IV bioreactor group (50 ± 3 μg/mg) was higher than of the subcutaneous group (26 ± 3 μg/mg) (*p* < 0.05), and similar to *in vitro* group 53 ± 2 μg/mg) (*p* < 0.05) at 3 weeks. These results support the ability of the IV bioreactor to maintain chondrogenesis of FCPC pellets observed above by biochemical analysis.

**Figure 5.**
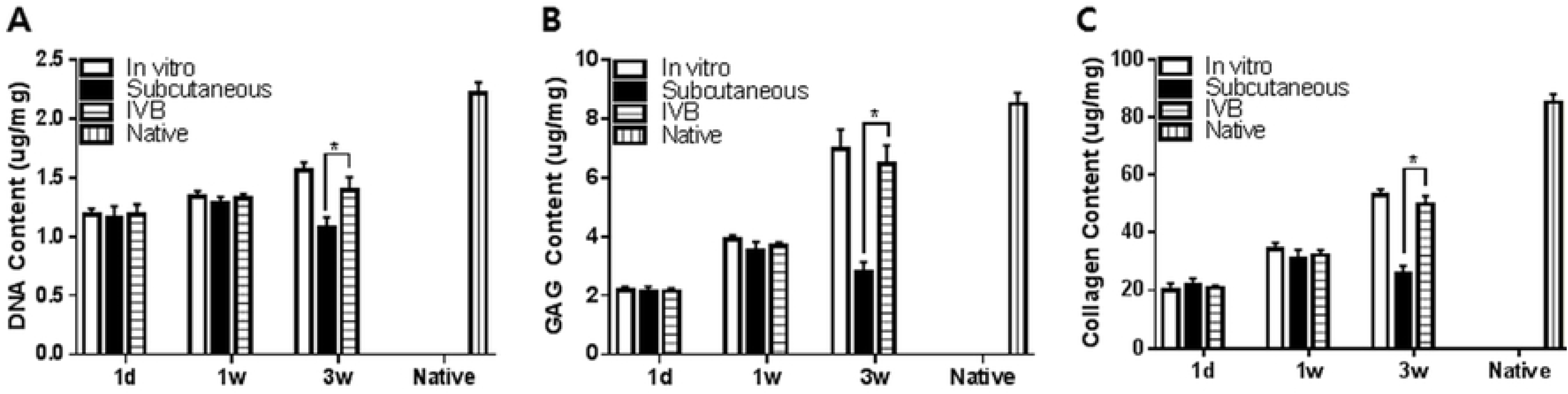
Biochemical analysis of *in vitro*, subcutaneous, IV bioreactor pellets and native rabbit cartilage (vertical striped bar) for total content of DNA (A), GAGs (B) and collagen (C). The pellets were cultured for 3 days *in vitro* before implantation in the subcutaneous and IV BIOREATOR of rabbit. The implanted pellets (*n* = 5/time point/group) were retrieved at 1day, 1 and 3 weeks after implantation. The amount of each component was calculated using standard curves for each assay and normalized by the wet weight of each pellet. Data are presented as a mean ± SD from 5 independent experiments in the histograms (*n* = 5). \**p* < 0.05

### 3.8. Molecular analysis of immunogenicity

The RT-PCR analysis showed that the mRNA of HLA-ABC was maintained significantly low levels in IV bioreactor group than in the *in vitro* group at all time points, and it decreased with time in all groups (Fig. 6A). The mRNA of CD80 was low levels only at 1 day and 1 week but increased at 3 weeks in all groups. However, its level of the IV Bioreactor was significantly lower than *in vitro* and subcutaneous groups. The mRNA pattern of CD86 showed a similar to that of the CD80. The mRNA of CD86 was no expression only at 1 day and 1 week but increased at 3 weeks *in vitro* and subcutaneous groups. However, its level of the IV bioreactor has maintained no expression at all time points. The western blotting analysis of protein showed a similar pattern overall to that of the mRNA (Fig. 6B). The protein of HLA-ABC *in vitro* group showed increase at 1 day and 1 week, and decrease at 3 weeks. But IV bioreactor and subcutaneous groups showed relatively low levels at 1 day, and it no expression at 1 and 3 weeks. The protein of CD80 was no expression only at 1 day and 1 week but increased at 3 weeks in all groups. However, its level of the IV bioreactor was significantly lower than *in vitro* and subcutaneous groups. The protein pattern of CD86 showed a similar to that of the CD80. The protein of CD86 was no expression only at 1 day and 1 week but increased at 3 weeks *in vitro* and subcutaneous groups. However, its level of the IV Bioreactor was maintained no expression at all time points. These results suggest that the IV Bioreactor not only better maintain the chondrogenic phenotype of FCPC pellets but also decreased the immunogenicity at a longer time when compared with the *in vitro* and subcutaneous.

**Figure 6.**
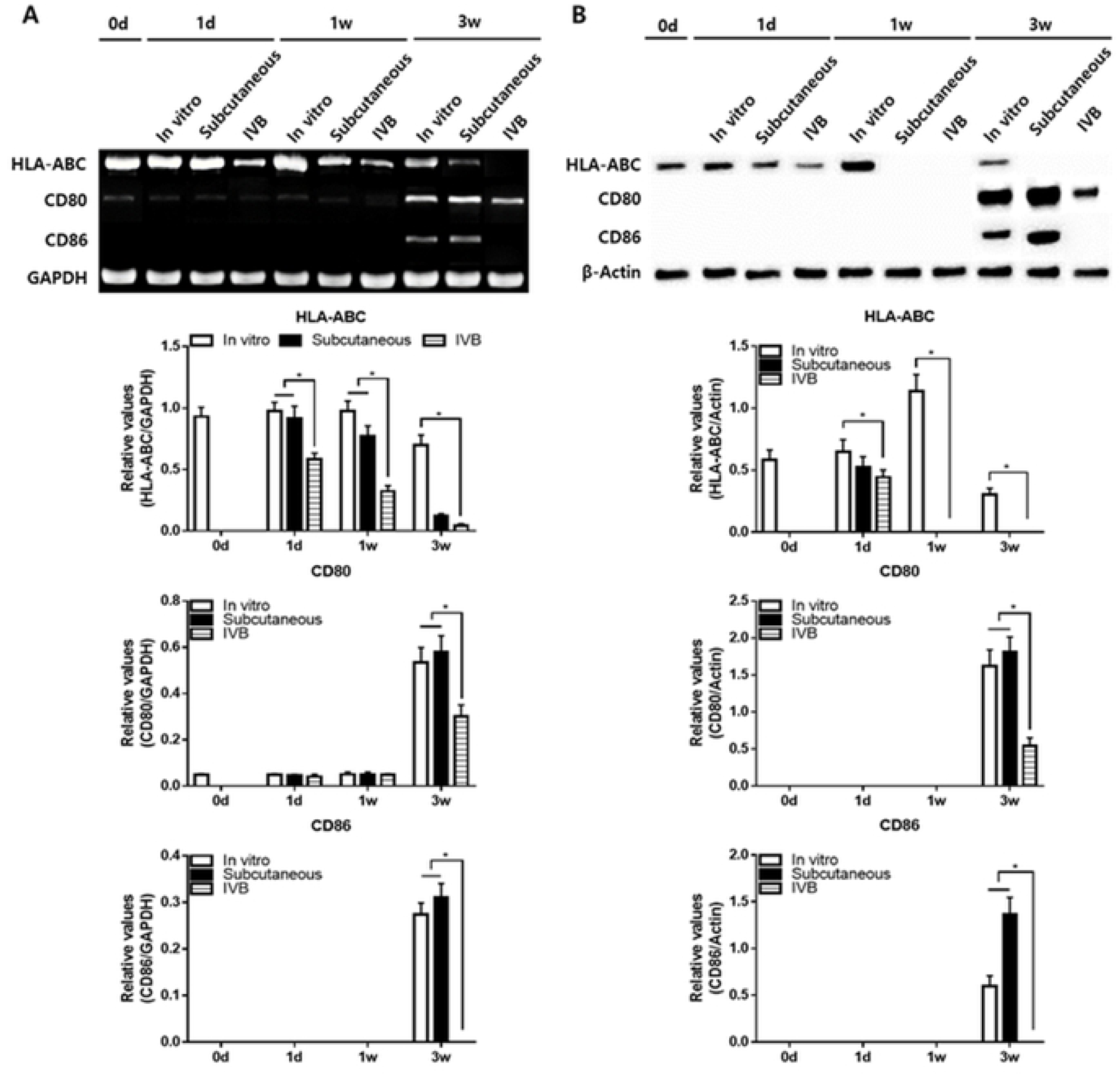
(A) Total RNA was isolated from the pellets and expression levels of HLA-ABC, CD80, CD86 and GAPDH were examined in the *in vitro*, subcutaneous and IV Bioreactor groups by RT-PCR analysis. Band intensities from 5 independent pellets (*n* = 5) were measured by the Image J program to obtain quantitative data. The intensities of HLA-ABC, CD80 and CD86 were normalized to that of GAPDH (a house-keeping gene) in the histograms. (B) Protein was isolated from the pellets and expression levels of HLA-ABC, CD80, CD86 and β-Actin were examined in the *in vitro*, subcutaneous and IV bioreactor groups by Western blotting. Band intensities from 5 independent pellets (*n* = 5) were measured by the Image J program to obtain quantitative data. The intensities of HLA-ABC, CD80 and CD86 were normalized to that of actin (a house-keeping gene) in the histograms.

### 3.9. Macroscopic and histological observation of the ectopic engineered cartilage for cartilage repair in vivo

The ectopic engineered cartilage was then applied to treat cartilage defect in the rabbit model. There was no infection, wound dehiscence, disability, and death throughout the *in vivo* experiment. The new tissue gradually formed and filled the defect from 4 to 12 weeks (Fig. 7A). In the control group, fiber-like tissue was gradually formed from 4 to 12 weeks. But the formation of new tissue was scarce and the defect center still has obvious depression at 12 weeks. Semi-transparent tissue gradually filled with defects *in vitro*, subcutaneous and IV bioreactor groups at 4 to 12 weeks. However, at 8 and 12 weeks, the similarity between neocartilage and native cartilage in the IV bioreactor group was better than that subcutaneous and *in vitro* groups. Specifically, the IV bioreactor group was almost completely covered with cartilage-like tissue, similar to the surrounding native cartilage with indistinct margins, after 12 weeks. But for the subcutaneous and *in vitro* groups, the margins between the native cartilage and the neocartilage was still obvious. The histological examination with safranin-O staining is shown (Fig. 7B). In the control group, fiber-like tissue formed in the defect, the integration between newly formed tissue and adjacent native cartilage can be easily identified with an obvious gap at 4 to 12 weeks. The thickness of neocartilage is similar to adjacent native cartilage and the staining intensity gradually increases in IV bioreactor, subcutaneous and *in vitro* groups at 4 to 12 weeks. In the IV bioreactor and *in vitro* groups, the neocartilage gradually became smooth of the surface, the cartilage lacuna density and distribution were similar to native cartilage, and the integration with adjacent native cartilage was complete. The IV bioreactor group at 12 weeks was better than *in vitro* group. However, in the subcutaneous group, the neocartilage surface was not smooth, the staining intensity and cartilage lacuna density were lower, and the integration with adjacent native cartilage was not complete. Consistently with the results of safranin-O staining examination, there was a statistically significant difference in the O’Driscoll score between the IV bioreactor, subcutaneous and *in vitro* group at each time point.

**Figure 7.**
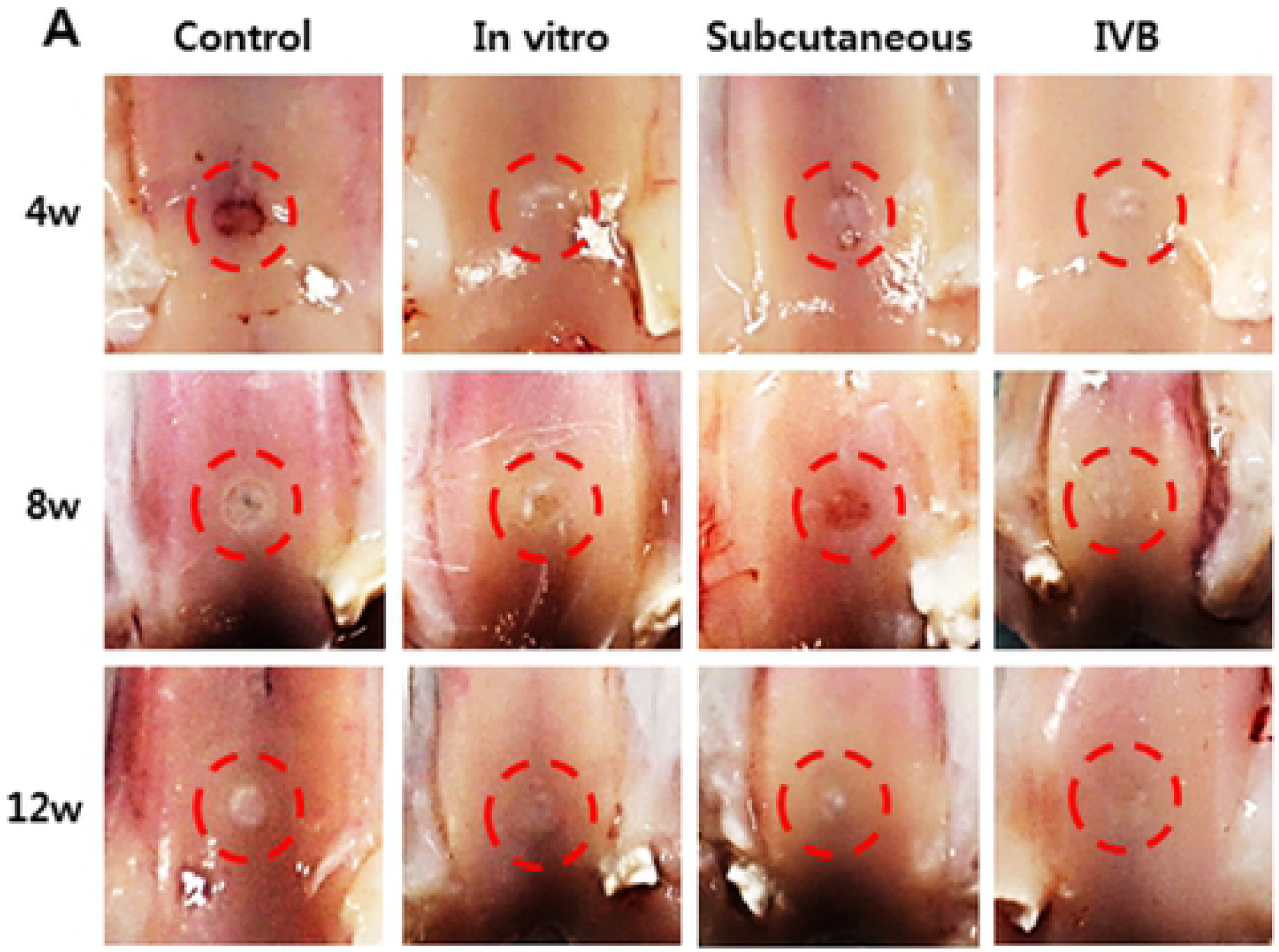
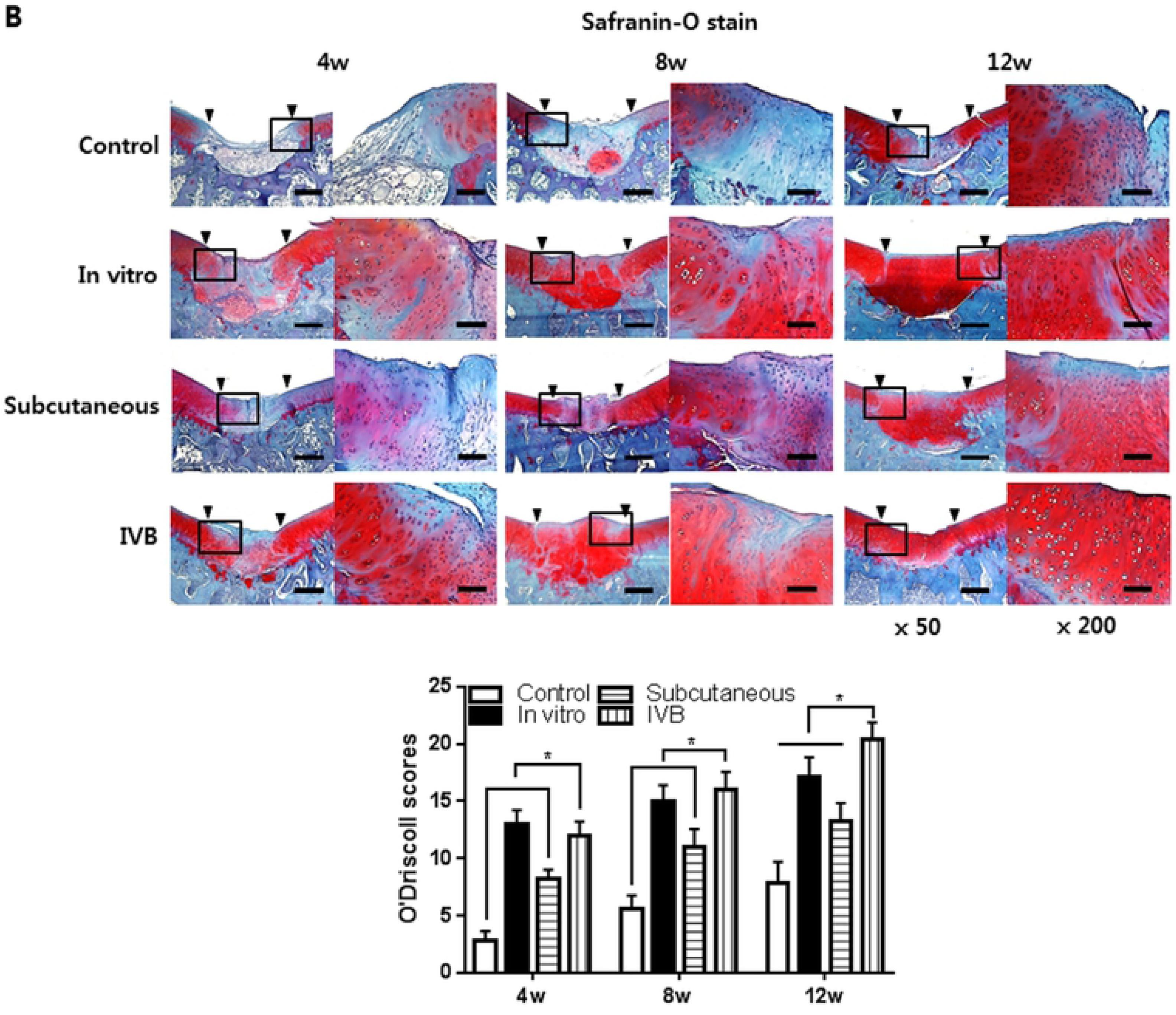
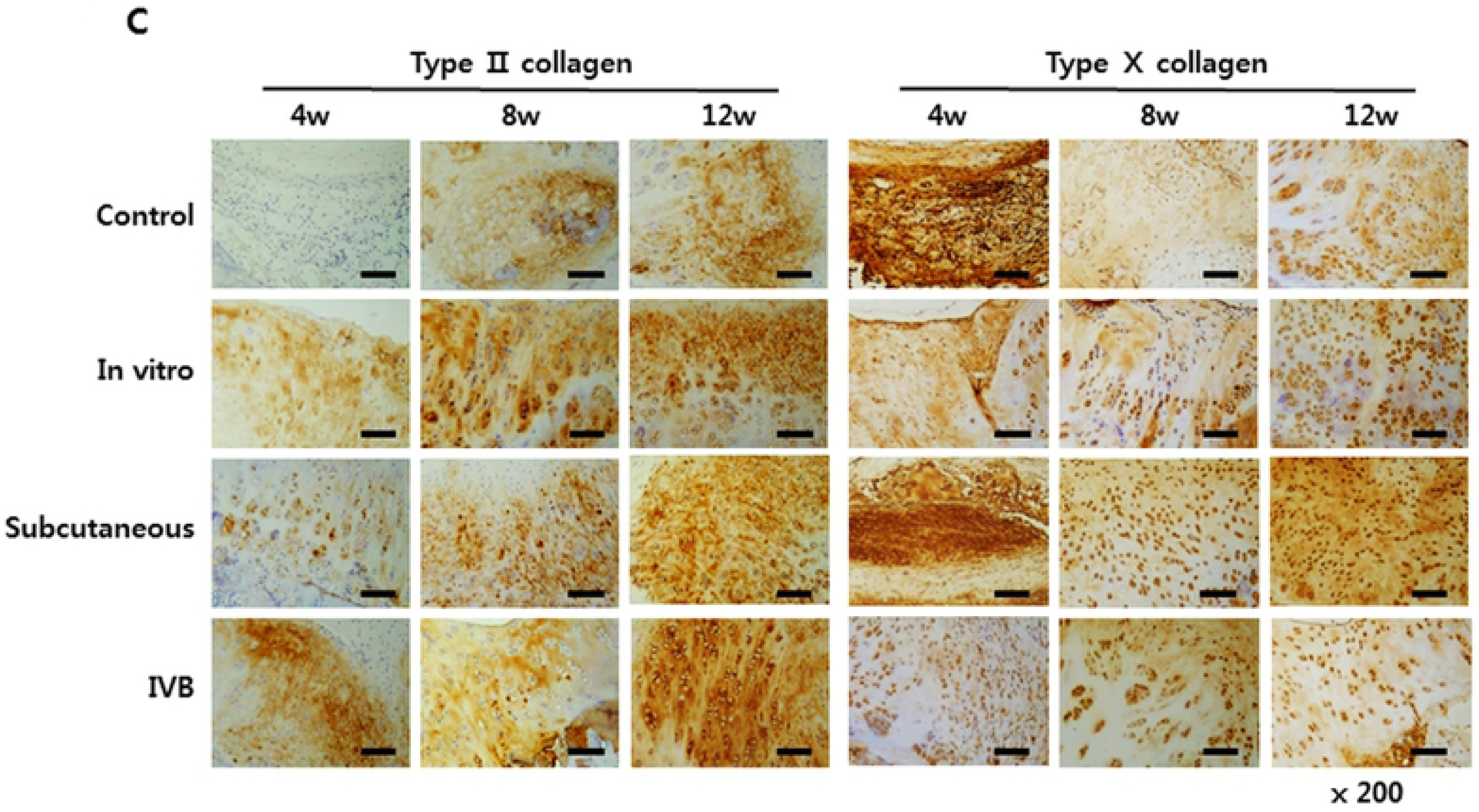
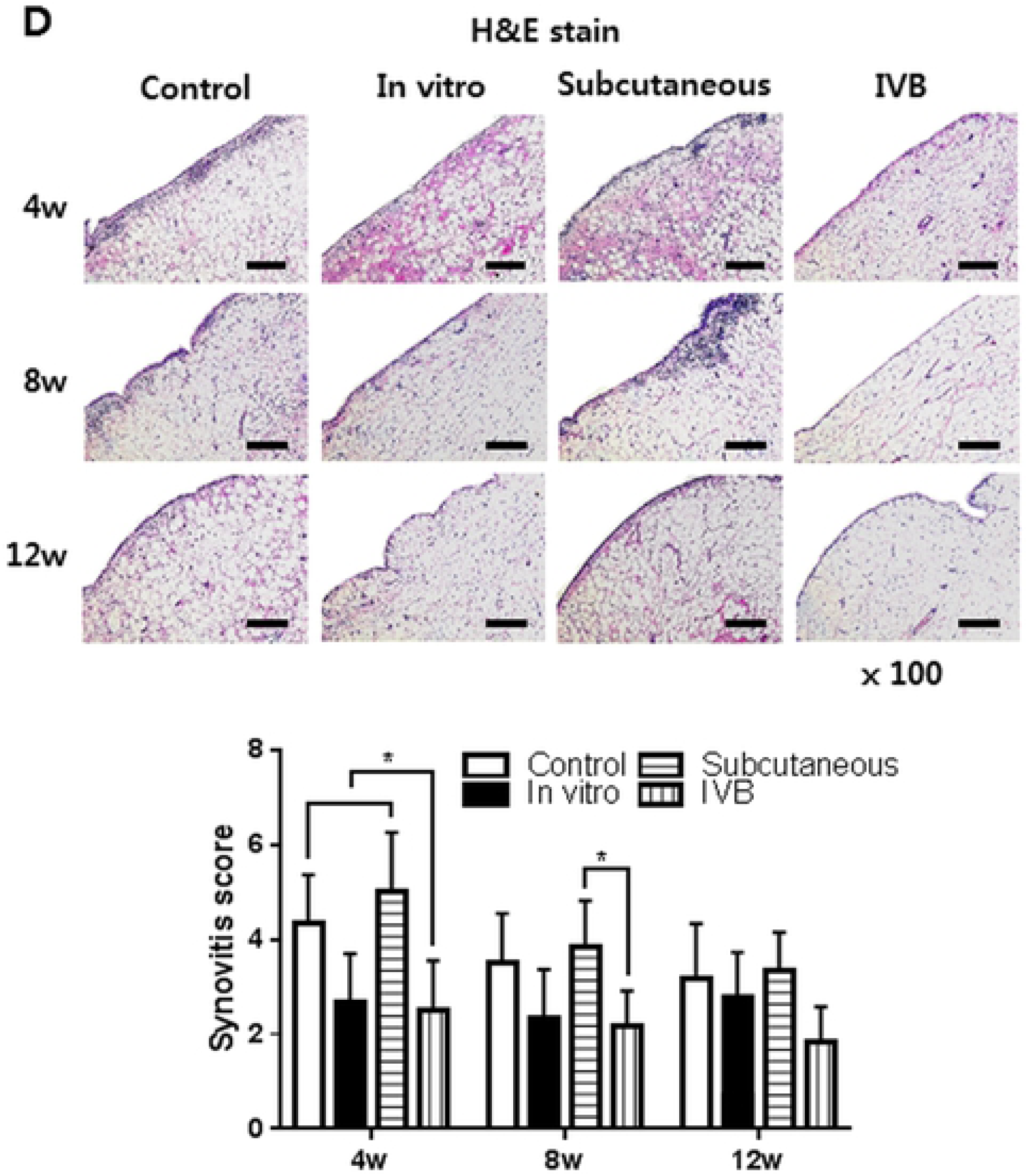
Macroscopic, histological and immunohistochemical observation of neocartilage in the cartilage defect at 4, 8 and 12 weeks post-implantation of ectopic engineered cartilage are shown. (A) Macroscopic observation of the newly formed tissue. (B) Safranin-O and fast green staining of neocartilage *in vivo*. And O’Driscoll scores were evaluated. (C) Immunohistochemical staining of type Ⅱ and type Ⅹ collagen. (D) H&E staining of synovium in the cartilage defect joint. And synovitis evaluated by Krenn’s synovitis scoring system. Scale bar = 400 μm for × 50, 200 μm for × 100 and 100 μm for × 200 images. Data are presented as a mean ± SD from 5 independent experiments in the histograms (*n* = 5). \**p* < 0.05.

### 3.10. Immunohistochemical observation of the ectopic engineered cartilage for cartilage repair in vivo

IHC staining of type Ⅱ and type Ⅹ collagen was performed to confirm the chondrogenic phenotype of implanted ectopic engineered cartilage and to evaluate the composition of newly formed tissues (Fig. 7C). As shown in the figure, the staining of type Ⅱ collagen in the IV bioreactor and *in vitro* group was stronger and uniform than the subcutaneous and control group, while the staining of type Ⅹ was weaker at the same time point. This suggests that the neocartilage of IV bioreactor group was close to hyaline cartilage. These results indicating that IV bioreactor can help ectopic engineered cartilage maintain better chondrogenic phenotype during cartilage repair *in vivo*.

### 3.11. Histological observation of the synovium

At 4 weeks, HE staining of the synovium in the subcutaneous and control groups showed an enlargement of the synovial lining cell layer, inflammatory cell infiltration and an increase of the cellularity, but gradually decreased with time (Fig. 7D). The synovium of the IV Bioreactor and *in vitro* groups were basically normal at each time point. Krenn’s synovitis score showed that there was a statistical difference between the IV bioreactor group and the subcutaneous group at 4 and 8 weeks, and no difference between the groups at 12 weeks. These results indicating that IV bioreactor not only helps ectopic engineered cartilage maintain better chondrogenic phenotype during cartilage repair *in vivo* but also may help reduce synovitis.

## 4. Discussion

From previous reports, ectopic cartilage reconstruction was not successful in immunocompetent animals, which mainly manifested as host immune response to allogenic or xenogeneic implant and cells, vascular invasion that may cause cartilage degradation and bone formation, host tissue ingrowth [18]. The results demonstrated that IV bioreactor is a useful experimental system for reconstruction of cartilage in immunocompetent hosts [19–21]. The IV bioreactor culture technique has been widely applied mainly to investigate cell differentiation. The pore size of cellulose membrane can be passed through most nutrients, but the cells cannot. The cellulose is a hydrophilic material, non-specific adsorption of proteins is low, mammalian cells are not easy to adsorb cellulose surface, so as to avoid the blocking of pores [22–25].

Recently, several studies demonstrated that nutrient supply of natural articular cartilage mainly comes from synovial fluid [26]. Our study shows that, IV bioreactor fluid component analysis showed that protein, glucose, and HA were close to synovial fluid than serum (Table 2). Glucose plays an important role in chondrogenesis and maintenance as a nutrient [27]. when HA was added to the culture medium, increase DNA, sulfated glycosaminoglycan and type Ⅱ collagen synthesis [28]. IV bioreactor environment promotes the synthesis of cartilage matrix and maintains the cartilage phenotype until 3 weeks, while subcutaneous cultured pellet degenerate and reduce the phenotypes after 3 weeks (Fig 4, 5). It may be because the IV bioreactor fluid contains substances required for chondrogenesis. IV bioreactor environment not only promotes the synthesis of cartilage matrix and maintains cartilage phenotype, but also delays the occurrence of calcification compared with subcutaneous (Fig. 4D). In this study, calcification in the subcutaneous group may be associated with host cell ingrowth and vascular invasion [29], while the reasons for delayed calcification in the IV bioreactor group may be related to some serum protein contained in the IV bioreactor, which may play an inhibitory role in the early stage of the calcification process after penetrating the matrix [30]. For another reason, although vascular invasion was avoided, osteogenic differentiation could not be completely prevented. It was inevitable that factors that would direct osteogenesis may be transmitted through body fluid during implantation. IVB environment not only promotes the synthesis of cartilage matrix and maintains cartilage phenotype, but also delays the occurrence of calcification compared with subcutaneous (Fig. 4D). Calcification is a multifactorial process caused by an imbalance between inhibitors and promineralization factors. Loss of proteoglycans often leads to articular cartilage calcification, because proteoglycans are effective mineralizing inhibitors [31]. The loss of chondrogenic phenotypes appears to be associated with increased matrix calcification (alizarin red staining) and hypertrophy (expression of type I and type X collagen). Chondrocyte phenotypic changes include hypertrophy differentiation, apoptosis, altered responses to growth factors, inflammatory cytokines and mediators [32]. Type I and type X collagen are related genes to the induction of chondrocyte hypertrophy [33].

In this study, RT-PCR and Western Blot results showed that the expression levels of HLA-ABC, CD80 and CD86 in pellets cultured for 1 week were lower, especially in IVB group than *in vitro* group and subcutaneous group (Fig. 6). HLA-ABC is a major histocompatibility complex class І (MHC І), CD80 and CD86 are co-stimulatory molecules. They are present on the cell surface and activate t cells when they binding to the t cell receptor (TCRs), initiating an immune response to destroy the foreign material [34]. Therefore, the presence of HLA-ABC, CD80, and CD86 on donor cells are important constituents involved in the immune rejection of implanted tissue engineered constructs [35]. Many researchers believe that the extracellular matrix (ECM) shield the MHC molecules from recognition by host cells; thereby protecting the chondrocytes from host immune responses [36]. In conclusion, this study successfully reconstructed cartilage in an immunocompetent host using a IV bioreactor system. The result revealed that an allogenous stem cell might also be applicable, which would resolve the problem caused by the shortage of resources for cartilage repair.

Compared with the *in vitro* group and subcutaneous group, IV bioreactor group healing the cartilage defects better, which was confirmed by Macroscopic, histological and Immunohistochemical observation (Fig. 7). The repair of cartilage defects using engineered cartilage in a IV bioreactor in vivo showed that neocartilage has formed around original cartilage and have a favorable boundary, whereas the constructs without the in vivo culture showed much less performance. This implied the superiority of in vivo engineered cartilage for repairing cartilage defects. According to previous reports, ectopic cartilage reconstruction has not been successful in immunocompetent animals, which is mainly manifested as host immune response to allogenic or xenogeneic implant and cells [18]. To prevent an immunological rejection, the scaffold and cells are encapsulated with the diffusion chamber before being implanted into the cartilage defect [37]. This may be because the IV bioreactor act as a mechanical barrier between the cell and the host, thereby hiding the surface antigen [19–21]. The results of synovium H&E staining and synovitis scoring showed that IV bioreactor group had lower synovitis scoring than the IV bioreactor group and subcutaneous group (Fig. 7D). The synovium of the subcutaneous group and the control group showed an enlargement of the synovial lining cell layer, cellularity increased and inflammatory cell infiltration. It is well known that the immune response of the joint to implanted neocartilage is mediated by the synovial membrane. Defects in articular cartilage can trigger an innate immune response even in the absence of an implant [38]. In this study, IV bioreactor was used to protect the pellet from host interference and create a good environment for cartilage regeneration.

## 5. Conclusion

In conclusion, this study successfully reconstructed ectopic cartilage in an immunocompetent host using a IV bioreactor system and obtaining satisfiable results applying for initial repair of cartilage defect. In addition, it was speculated that, because the reconstructed tissue to repair the defect of host was formed right in host, it might be preferred to extracted allografts.

## Funding

This paper was funded by the Ministry of Health and Welfare (HI17C2191) in 2017.

## Competing interests

There is NO Competing Interest.

## Author contributions

**Xue Guang Li**: Conception and design, collection and assembly of data, data analysis and interpretation, manuscript writing

**In-Su Park**: Conception and design, collection and assembly of data, data analysis and interpretation, manuscript writing

**Byung Hyune Choi**: Conception and design, collection and assembly of data, data analysis and interpretation

**Ung-Jin Kim**: data analysis and interpretation

**Byoung-Hyun Min**: Conception and design, financial support, manuscript writing, final approval of the version to be submitted

